# A versatile cryopreservation method for peri-gastrulation squamate embryos optimised using the veiled chameleon (*C. calyptratus*)

**DOI:** 10.64898/2026.04.01.715795

**Authors:** A Weberling, M Durnin, NA Shylo, MC McKinney, H Wilson, R Kupronis, SA Williams, PA Trainor

**Affiliations:** All Souls College, University of Oxford, UK; Nuffield Department of Women’s and Reproductive Health; Stowers Institute for Medical Research, Kansas City, MO, United States; Department of Biological and Biomedical Sciences, Rowan University, Glassboro, NJ, United States; Department of Anatomy and Cell Biology, University of Kansas Medical Center, Kansas City, MO, United States

## Abstract

Stem cell technologies have become a vital component of conservation efforts around the globe. Biobanks and pluripotent stem cell lines help to ensure species and their genetic diversity are preserved. These efforts have however, focussed mostly on mammals and birds, and the cryopreservation protocols for embryos and cells were developed decades ago laying the basis for artificial reproductive techniques for species conservation. With over 20% of non-avian reptile species facing extinction, it is imperative to establish protocols for reptiles to ensure species preservation and also to facilitate the establishment of new reptile model organisms to match the standard of mammals. Here, we have generated a cryopreservation method for preserving early gastrulating veiled chameleon embryos as a representative squamate species. To this end, we first developed a tissue culture method for maintaining cells extracted from peri-gastrulation chameleon embryos and then tested different cryopreservation methods altering the concentration of the penetrating cryoprotectant DMSO and assessing the effect of the addition of non-penetrating cryoprotectants Trehalose and Sucrose. We then optimised a protocol for whole embryo vitrification in 20% DMSO with added Trehalose or Sucrose that can easily be adapted for fieldwork. Taken together, our method not only provides a protocol for conservation efforts but also lays the basis for mechanistic studies of early squamate embryo development by enabling cryopreservation of whole embryos in a fieldwork setting, which facilitates their live transport back to a laboratory for functional experiments or molecular analyses.

## Introduction

In the face of global extinction, the risk for mammals, birds and amphibians has been tracked continuously, however the first assessment for non-avian reptiles was only carried out in 2022. This study on reptile extinction risk focussed on squamates, turtles, crocodiles, and tuataras and reported 21.1% of reptile species as being threatened (Cox et al., 2022). Next to traditional conservation methods that include habitat preservation and dedicated breeding schemes, assisted reproductive technologies (ARTs) that comprise biobanking of germplasm, embryos and tissues as well as the generation of pluripotent stem cells, *in vitro* gametogenesis, artificial insemination, and *in vitro* fertilisation are each emerging as vital components of conservation science (Hutchinson et al., 2024). A focal point of all these techniques is the controlled preservation and storage of the respective tissues through cryopreservation.

Cryopreservation enables freezing, unlimited storage in liquid nitrogen and thawing of a specimen while it remains alive. First pioneered for sperm (Polge et al., 1949), it has since become a vital tool for ARTs, and preserving sperm, oocytes, zygotes and blastocysts have greatly enhanced the success rates of ARTs (Casciani et al., 2023). In addition to worldwide interest in fertility preservation for humans, cryopreservation also emerged as a vital tool in conservation science for biobanking tissues of endangered species (Chen and Mastromonaco, 2025; Hutchinson et al., 2024) as well as in the fields of developmental and stem cell biology, by facilitating the storage of stem cell lines. Cryopreservation is generally carried out in a specific medium that contains a penetrating cryoprotectant such as DMSO, ethylene glycol, 1,2-propandiol, and glycerol (Pegg, 2007) which may be combined with a non-penetrating cryoprotectant such as Sucrose or Trehalose or high molecular weight polymers (Yong et al., 2020). Consequently, a large variety of combinations of cryoprotectants and cryopreservation techniques exist, which have been optimised for different tissue types and species (Bartolac et al., 2018; Sanaei et al., 2025). Cryopreservation follows one of two major freezing protocols. For slow-freezing, the specimen is cooled stepwise with the consecutive dehydration of the cells preventing ice-crystal formation. This method is often used for cell suspensions and the freezing of large tissues (Khaydukova et al., 2024). During vitrification, the specimen is cooled rapidly which also prevents ice-crystal formation resulting into the specimen adopting a glass-like form. Vitrification is the preferred method of gamete and early embryo preservation (Mandawala et al., 2016). While cryopreservation methods were developed for many live-stock embryos and tissues (Dobrinsky, 2002) and successful efforts have been made for numerous mammalian species (Silva et al., 2015), no studies have focussed on the preservation of non-avian reptile embryos other than the development of cryopreservation protocols for sperm (Banchi et al., 2025; Campbell et al., 2020; Sánchez-Rivera et al., 2022; Young et al., 2022).

Squamates, which include all lizards, worm lizards, and snakes, constitute the largest order of non-avian reptiles with over 12,000 species described to-date (Uetz and Hošek, 2025). Having branched from other reptiles about 240MYA (Gable et al., 2023), squamates spread across the entire globe and exhibit highly diverse phenotypes as adaptations to their respective ecological niches (Roll et al., 2017). Despite the abundance of squamates, few species have emerged as model organisms with none of the conventional model organisms in biology being a squamate. Thus, many standard tools in developmental biology are not available for squamates, or have only recently been adapted for this group. In the past decade, several molecular tools such as CRISPR/Cas mediated gene editing, *in vitro* embryo culture, and live imaging, as well as cell culture were pioneered for different squamate species (Rasys et al., 2019; Samudra et al., 2024; Shylo et al., 2023).

The veiled chameleon (*C. calyptratus*) is a representative of the squamate suborder Iguania. In contrast to many squamates where pre-oviposition development continues up to the early organogenesis and limb bud formation stages, the veiled chameleon initiates gastrulation upon oviposition (Diaz Jr et al., 2019; Peter, 1934; Weberling et al., 2025). In addition, while other squamate model species lay only 1 egg at a time such as members of the genus Anolis (Andrews and Rand, 1974) or 1-2 eggs per clutch such as members of the genus Gekkonoida (Kluge, 1987), veiled chameleon females can lay clutches of 45-90 roughly time-matched eggs (Diaz et al., 2015). The early embryonic stage of development at oviposition together with large clutch sizes, makes the veiled chameleon an ideal model to develop a cryopreservation protocol for peri-gastrulation squamate embryos.

Here, we developed a versatile method to cryo-preserve peri-gastrulation veiled chameleon embryos. We first established a tissue culture system for maintaining cells extracted from chameleon embryos for 48h and then carried out cryopreservation trials and optimisation to develop a cryopreservation protocol that ensures high rates of cell survival. This is – to our knowledge – the first cryopreservation protocol for non-avian reptile embryos, which will be highly impactful for the emerging field of early reptile developmental biology, and will also prove extremely useful for conservation purposes. Our method provides a straightforward setup which can be adapted to field work in any setting, enabling the collection of live embryos for functional experiments and stem cell derivation in a lab setting or for biobanking of early embryos for primordial germ cell isolation for species conservation.

## Results

### Veiled chameleon embryos can be cryopreserved at 0dpo

Upon oviposition, the veiled chameleon embryo has a compact, round shape (Figure 1A/B) (Peter, 1935; Weberling et al., 2025). The embryonic epiblast (e) has folded to give rise to the amniotic cavity (ac), which is formed on the dorsal side by the epiblast-derived amnion-like tissue (a) with the epiblast on the ventral side (Figure 1A/C/D). Dorsally, the embryo is covered by the trophoblast-like tissue (t) that is formed by enlarged, double-nucleated cells (Figure 1C/Di) (Weberling et al., 2025), while the epiblast is ventrally enveloped by the hypoblast (h) (Figure 1Cii). At 0 days post oviposition (dpo), the chameleon embryo has completed anterior-posterior patterning and initiated gastrulation visible by mesoderm formation at the posterior side (m), which is distinguishable by a thickening of the epiblast layer (Figure 1Di). The site of gastrulation can be distinguished through mesoderm ingression (Figure 1E arrows) with the mesoderm migrating along the epiblast (asterisks). Taken together, the veiled chameleon comprises 5 major tissues at 0dpo - epiblast, trophoblast-like, hypoblast, amnion-like, and mesoderm - which could potentially be isolated to generate primed pluripotent chameleon embryonic stem cells, as well as trophoblast, hypoblast, and amnion cell lines.

**Figure 1:**
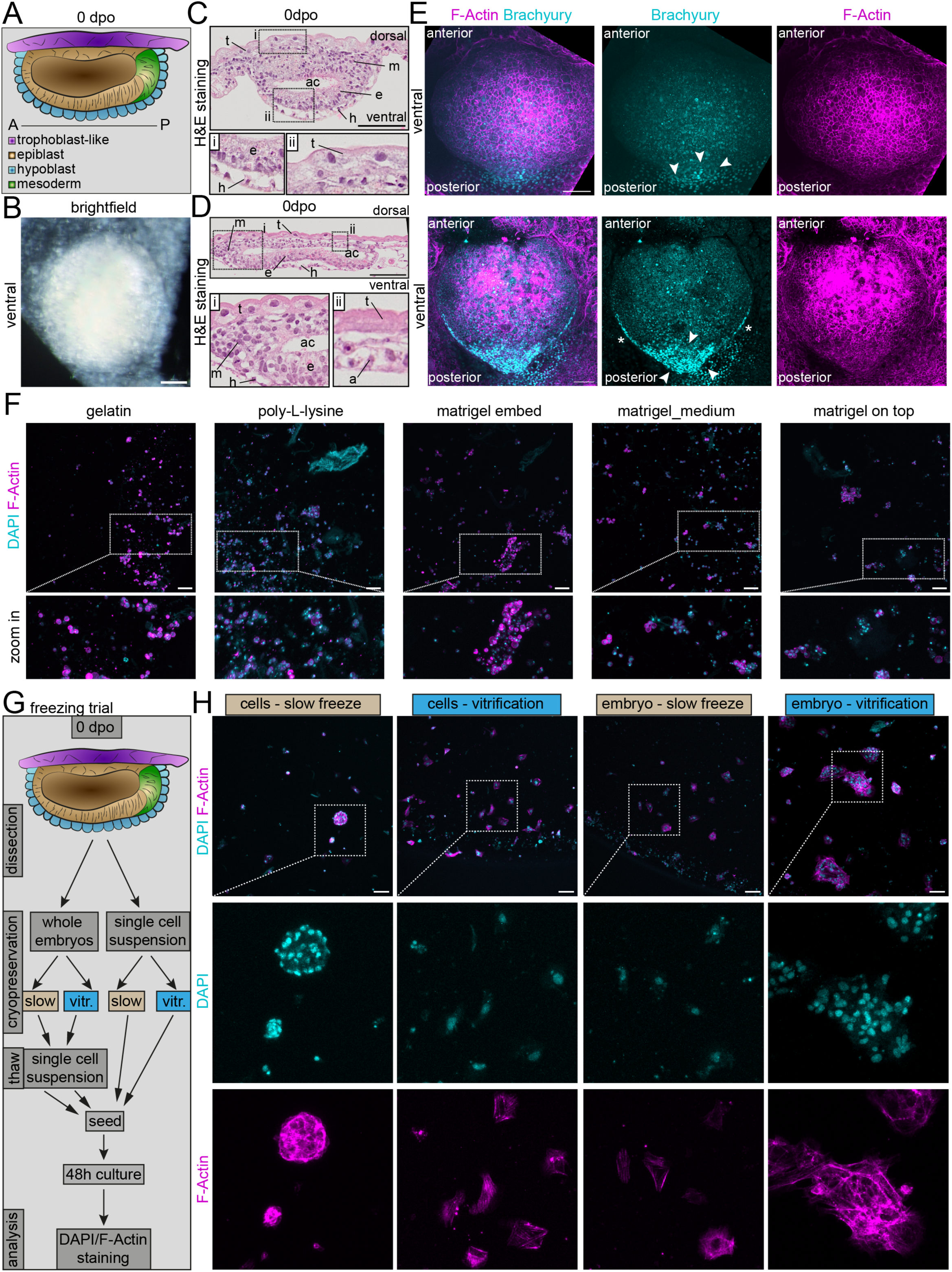
0dpo chameleon embryos can be cryopreserved. **A.** schematic drawing of a 0dpo chameleon embryo annotating the epiblast (beige), hypoblast (cyan), trophoblast-like tissue (magenta), and nascent mesoderm (green). **B.** brightfield image of a ventral view of a 0dpo embryo. **C/D.** Hematoxylin & Eosin (H&E) staining of cross sections of a 0dpo embryo. Boxes highlight higher magnification insets (i/ii). a=amnion, ac=amniotic cavity, e=epiblast, h=hypoblast, m=mesoderm, t=trophoblast-like tissue. **E.** Immunofluorescence staining of 0dpo embryo. F-actin (magenta) and Brachyury (cyan). Maximum intensity projection of the ventral side. Top row: site of gastrulation visible by brachyury-expression (arrows). Bottom row: clear symmetry breaking at the site of gastrulation (arrows). The mesoderm migrates along the sides of embryo (asterisks) **F.** Tissue culture trial. 0dpo embryos dissociated and cells cultured for 48h. Wells coated with gelatin (left), poly-L-lysine (second left) or cells embedded in matrigel (middle), cells seeded on top of matrigel, with matrigel in medium (second right), cells seeded with matrigel in medium (right). DAPI (cyan) and F-Actin (magenta). Representative of 3 biological replicates. Box indicates higher magnification inset. **G.** Schematic drawing of freezing trial set up. Embryos were either suspended as single cells following trypsinisation and then subjected to slow freezing or vitrification or alternatively frozen as whole embryos following slow freezing or vitrification protocols. Following thawing, the cells were seeded directly while embryos were trypsinised and then seeded. After 48h culture, the cells were fixed and analysed. **H.** Results of freezing trial. DAPI (cyan) and F-Actin (magenta) staining. Boxes indicate higher magnification insets. Representative images of 3 biological replicates images of replicates 2&3 in Figure S1BA. All scale bars 100um.

We decided to use cell culture of dissociated chameleon embryos as a readout to assess the success of cryopreservation since not all cells in an embryo might survive the freeze-thaw process. Also, different embryonic tissue types may have different post-thaw survival rates. Since no cell culture method for 0dpo chameleon embryonic, trophoblast, or hypoblast stem cell lines existed, we therefore initiated tissue culture trials with 0dpo embryos to establish the conditions that would allow for 48h cell culture survival post-thaw. For this, we dissociated 0dpo embryos using trypsin and seeded the near single cell suspension into differently coated wells using a culture medium that has been optimised for chameleon fibroblast culture (Shylo et al., 2024). As the cells did not attach to the plastic dish, we tested different coating (gelatin, poly-L-lysine) and embedding techniques (matrigel) (Figure 1F). While we could observe cell attachment and survival in all conditions, gelatin-coating appeared to result in less debris, and was therefore used in further development of the protocol.

Next, we set out to develop a cryopreservation method. For this, we first tested whether slow freezing or vitrification would provide better embryo survival and whether freezing of single cell suspensions or whole embryo freezing would yield better results. We therefore subjected whole embryos or single cells to vitrification or slow freeze cryopreservation (Figure 1G). Following thawing, the whole embryos were dissociated to a near single cell suspension and seeded. The cells originating from embryos dissociated prior to cryopreservation were centrifuged following thawing, the pellet resuspended in medium and then seeded. Following 48h of culture, the cells were fixed and stained for DAPI and F-actin to assess cell numbers and cell morphology. While we could observe live cells under all cryopreservation methods tested, the slow freezing of single cell suspensions yielded low numbers of surviving cells. Interestingly, the cells did not seem to adhere well to the dish and seemed quite round with the cytoskeleton directly arranged around the nucleus (Figure 1H/S1A, column 1). Vitrification of single cell suspensions also yielded low cell numbers but resulted in better attached cells. Here, F-actin staining revealed that the cells spread over the dish as originally expected since the culture medium does not provide any stemness-retaining factors (Figure 1G/S1A, column 2). Cells originating from whole embryos subjected to a slow freeze, and then single cell suspensions post thawing, were also observed to be attached to the culture dish (Figure 1G/S1A column 3). Cells seeded following whole embryo vitrification and single cell suspension, also attached well and qualitatively exhibited the best survival (Figure 1H/S1A column 4). Overall, both vitrification methods resulted in less debris in the wells. To understand whether adding matrigel to the medium would further enhance cell survival, we assessed the morphology of cells seeded from vitrified whole embryos (Figure S1B). As we could not observe a qualitative difference between gelatin-coated dishes and gelatin-coated dishes with matrigel in the medium, we decided to proceed with gelatin-coated dishes.

While single cell suspensions of embryos prior to freezing enable stem cell generation, and methods such as single cell RNA sequencing, the intact embryo is not maintained for spatial analysis or *in vitro* culture. As vitrification of whole embryos can be more easily achieved in a fieldwork setting than slow freezing and single cell suspension, (with the need to dissociate the embryos in an incubator), we decided to optimise the whole embryo vitrification protocol.

### Optimisation of a whole embryo vitrification protocol

To optimise our vitrification method, we varied the concentration of DMSO (10/20%) and added either Trehalose (0.25M) or Sucrose (0.5M) as further non-penetrating cryoprotectants. Following thawing, the embryos were dissociated and seeded into individual wells of a 96-well plate. After 48h culture, the cells were fixed and analysed (Figure 2A).

**Figure 2:**
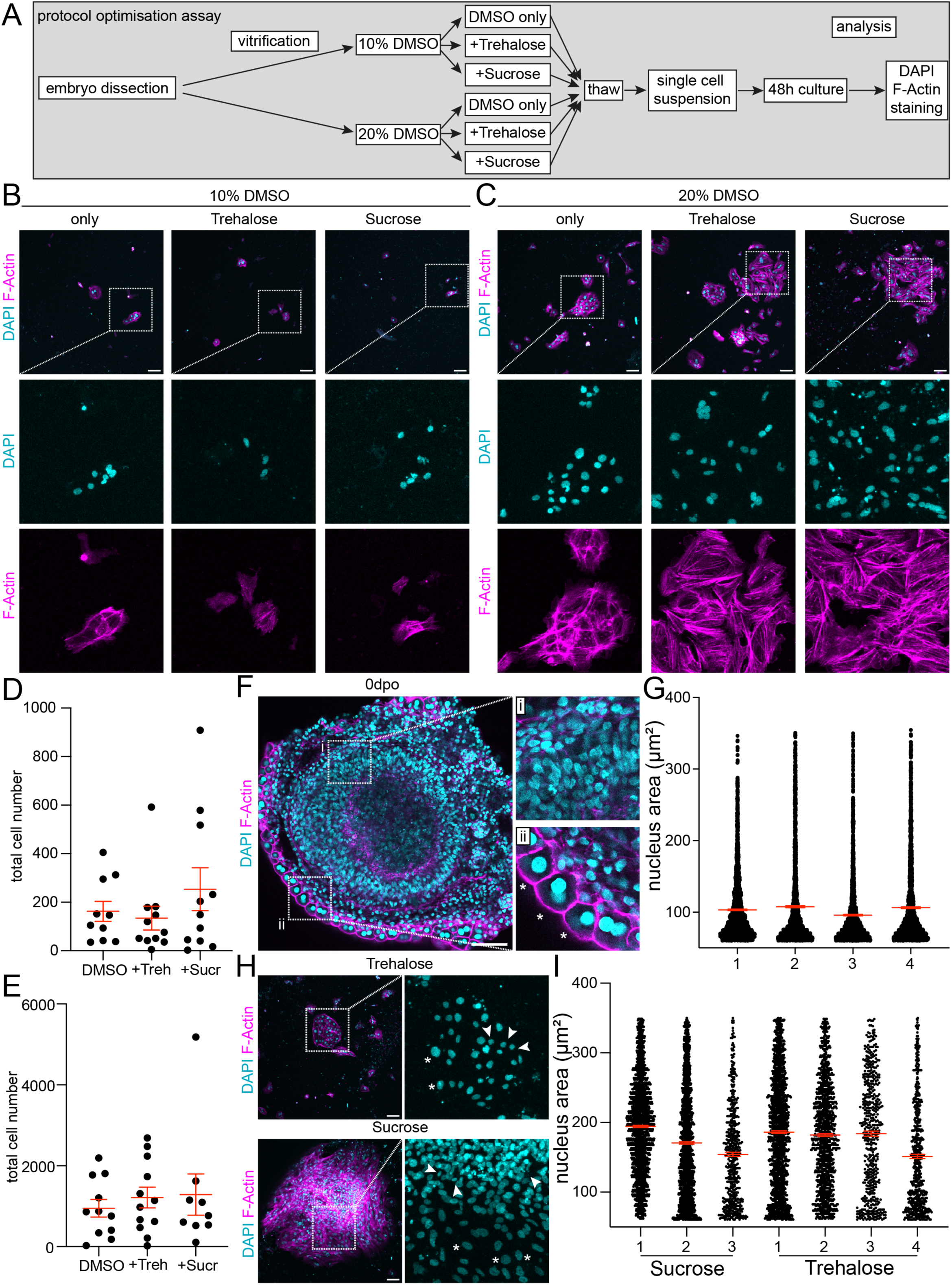
Optimisation of the cryopreservation protocol. **A.** Schematic of cryopreservation optimisation assay. **B/C.** Maximum intensity projections of cells cultured for 48h following thawing and single cell suspension. (B) Embryos vitrified in 10% DMSO (+/-) Trehalose/Sucrose. (C) Embryos vitrified in 20% DMSO (+/-) Trehalose/Sucrose. DAPI (cyan) and F-Actin (magenta) staining. Boxes indicate higher magnification insets. **D/E.** Quantitative analysis of the cryopreservation assay. Scatter plot ± SEM of cell numbers of embryos vitrified in 10% (D) or 20% (E) DMSO±Trehalose/Sucrose. **F.** Confocal imaging of a 0dpo embryo highlighting nuclear size differences in different tissues. Boxes indicate higher magnification insets. (i) Epiblast and hypoblast cells exhibit smaller nuclei. (ii) Trophoblast cells (asterisks) possess larger nuclei and are double nucleated. **G.** Quantitative analysis of nuclear area using the StarDist plugin. Area cut-offs of 60um-350um were used to exclude apoptotic cells and merged nuclei groups. Scatter plot of 4 individual embryos, mean ±SEM. **H.** maximum intensity projection of cells cultured for 48h following thawing and single cell suspension. Top row vitrification in 20% DMSO +Trehalose, bottom row vitrification in 20% DMSO+Sucrose. DAPI (cyan) and F-Actin (magenta staining. Boxes indicate higher magnification insets. Arrows indicate potential epiblast/hypoblast (small nuclei), asterisks highlight potential trophoblast-like cells (large nuclei). **I.** Quantitative analysis of nuclear area using the StarDist plugin. Area cut-offs of 60um-350um were used to exclude apoptotic cells and merged nuclei groups. Scatter plot of individual experiments, each experiment consisted of 3 individual embryos vitrified/seeded, mean ±SEM. All scale bars 100um.

We initially assessed the effect of 10% DMSO alone and then by adding Trehalose or Sucrose. 10%DMSO only, resulted in low density single cells or small groups of cells that attached and spread over the plastic similar to the cryopreservation trial (Figure 2B/S2A left). Addition of Trehalose or Sucrose resulted in a slight qualitative increase in cell numbers with less debris in the wells but did not exhibit a marked difference overall (Figure 2B/S2A middle/right). We then analysed the effect of increasing the DMSO concentration to 20% alone and then by adding Trehalose or Sucrose. 20% DMSO only, resulted in markedly higher cell numbers (Figure 2C/Figure S2B left). However, after adding either Trehalose or Sucrose, we observed a massive comparable increase in cell number (Figure 2C, S2B middle/right), demonstrating that these two conditions both appeared to provide a similar protective effect. To gain further insight into the effect of 10% versus 20% DMSO and the addition of Trehalose or Sucrose, we carried out a quantitative analysis of total cell number (Figure 2D/E). This confirmed our observation that vitrification in a freezing medium containing 20%DMSO +Trehalose/Sucrose resulted in much higher cell numbers from a mean of 162 cells in 10%DMSO only conditions, to a mean of 1288 cells per well in 20%DMSO+Sucrose media. Our optimisation assay showed that vitrification in 20%DMSO+Trehalose or Sucrose was a far superior cryopreservation method for chameleon embryos than 10% DMSO only.

Next, we sought to gain a deeper understanding of the cell composition of the cells surviving vitrification. Although we achieved high cell survival rates using our optimised cryopreservation protocol, we did not have any information about whether all tissues were preserved at the same rate. Since this protocol is designed for full embryo preservation as well as a means to generate embryonic stem cells, it is important that all cell types survive. Unfortunately, we have not yet established markers similar to those used to define specific tissue types such as Oct4 and Sox2 for the epiblast, or Cdx2 for the trophoblast, and Gata6 for the hypoblast in mouse embryos (Molè et al., 2020). Commonly used antibodies have also not worked because they are either not specific for chameleons or because the proteins are not expressed in these tissues (Weberling et al., 2025). However, we did observe that different tissues exhibit cells with differently sized nuclei (Figure 2F). The nuclei of epiblast and hypoblast cells are much smaller (Figure 2Fi) than the nuclei of the trophoblast-like tissue, the cells of which also exhibit double-nucleation (Figure 2Fii). While the mechanism underlying these different tissue-specific characteristics still need to be elucidated (Weberling et al., 2025), they may be used as a proxy to distinguish different tissue types. We used an automated nuclear segmentation program to measure nuclear areas in 0dpo embryos (Schmidt et al., 2018), and assessed nuclear areas in 4 individual embryos (Figure 2G). Each embryo exhibits a wide range of nuclear areas with an average mean of about 100um^2^. We then analysed the nuclear areas of cells cultured following cryopreservation with our two optimal methods (Figure 2H) and in both cases observed enlarged nuclei (arrows) and markedly smaller nuclei (asterisks). We carried out quantitative analysis and observed that both conditions captured the same range of nuclear sizes though the mean had shifted towards 150-200um^2^ (Figure 2I). While this points towards trophoblast-like cells being more successfully preserved than epiblast or hypoblast cells, it is still clear that we obtain cells of all nuclear sizes suggesting that cells of each tissue type survive cryopreservation with our optimised protocol.

### Optimised vitrification protocols result in better whole embryo survival

Following the optimisation of our cryopreservation protocol, we next aimed to understand whether the different survival rates may also be reflected in different embryo morphologies post-thawing. For this, we fixed embryos immediately after thawing and assessed overall morphology with specific attention to epiblast tissue integrity (Figure 3A). Embryos vitrified in 10% DMSO retained overall embryo integrity (Figure 3A left column), with the majority of the epiblast tissue preserved (zoom-in). However, lumen formation did not appear to be complete as the embryo exhibited large breaks (arrows). Similarly, hypoblast integrity was broken on the distal side of the embryo (asterisks). Embryos frozen in 10% DMSO/Trehalose appeared to have better preserved overall morphology (Figure 3A middle column) but still exhibited small breaks in the epiblast tissue (Figure 3A, middle column, arrow). The hypoblast however, appeared intact. Embryos frozen in 10% DMSO/Sucrose, appeared to have an intact epiblast and hypoblast (Figure 3A right column). When vitrified in 20% DMSO, the epiblast was fully intact but small breaks occurred in the hypoblast (Figure 3B left column, asterisks). 20% DMSO/Trehalose and 20% DMSO/Sucrose both yielded intact embryos that contained an intact epiblast and hypoblast (Figure 3B middle/right columns). The trophoblast lineage was also retained in each condition however, it curled up which prevented a proper assessment of tissue integrity (Figure 3A/B top row yellow outlines). Taken together, overall embryo integrity is maintained in all vitrification conditions.

**Figure 3:**
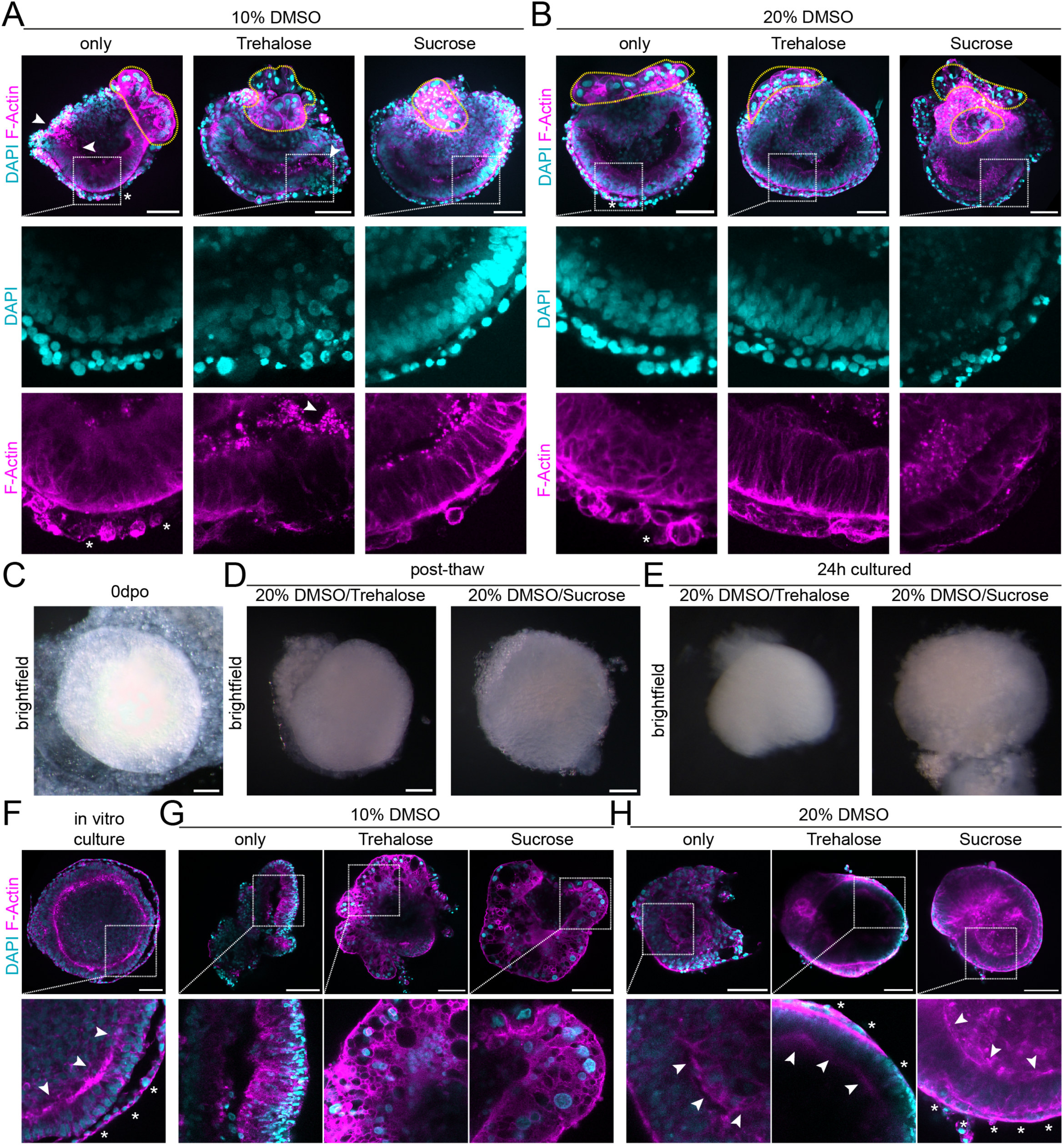
optimised vitrification method results in better whole embryo survival. **A/B.** Embryo morphology following cryopreservation in 10% DMSO (+/-) Trehalose/Sucrose (A) or 20% DMSO (+/-) Trehalose/Sucrose (B). Confocal imaging of DAPI (cyan) and F-Actin (magenta) staining of a virtual cross section. Boxes indicate higher magnification insets. Yellow outline indicates trophoblast-like tissue. Asterisks indicate hypoblast break points. Arrows indicate epiblast break points. **C.** Brightfield image of a 0dpo embryo from the ventral side. **D.** Brightfield image of embryos post thawing following vitrification in 20% DMSO/Trehalose (left) or 20% DMSO/Sucrose (right). **E.** Brightfield image of vitrified embryos following 24h of *in vitro* culture. Left, embryo vitrified in 20% DMSO/Trehalose, right, embryo vitrified in 20% DMSO/Sucrose. **F-H** Confocal images of embryos after *in vitro* culture. (F) 0dpo embryo cultured for 24h. (G) 0dpo embryo cultured for 24h following vitrification in 10% DMSO (+/-) Trehalose/Sucrose. (H) 0dpo embryo cultured for 24h following vitrification in 20% DMSO (+/-) Trehalose/Sucrose. DAPI (cyan) and F-Actin (magenta) staining. Boxes indicate high magnification insets. Arrows indicate epiblast. Asterisks indicate hypoblast. All scale bars 100um.

We then cultured whole post-thawing embryos to understand whether one or more of the vitrification conditions may lend itself to whole embryo survival. First, we carried out a brightfield assessment of 20%DMSO/Trehalose and 20% DMSO/Sucrose vitrified embryos post thawing and then following 24h culture. Embryos that had been preserved following other conditions such as 10% DMSO fell apart and as such we did not take brightfield images. At 0dpo, the embryo is a tight, solid structure (Figure 3C/S3A). Embryos assessed directly post-thawing exhibited high similarity to 0dpo embryos (Figure 3D/S3B). Following 24h culture, embryos stemming from vitrification in 20%DMSO/Trehalose or 20% DMSO/Sucrose remained intact overall but exhibited loose hypoblast or trophoblast structures (Figure 3E/S3C).

We then carried out more detailed, qualitative morphological assessments of embryos cultured post thawing. Since no tissue specific markers are available for these early-stage chameleon embryos, we based our analysis on morphology using DAPI/F-Actin staining. A 24h *in vitro* cultured 0dpo embryo exhibits a large epiblast, with an intact but possibly loose hypoblast (Figure 3F). Embryos vitrified using 10% DMSO (+/-) Trehalose/Sucrose do not retain an organised epiblast compartment and fall apart (Figure 3G). Embryos vitrified in 20% DMSO appeared to retain a very small epiblast compartment surrounding a central lumen within disorganised tissue (Figure 3H left, yellow outline). Interestingly, embryos vitrified in 20% DMSO Trehalose exhibited a full epiblast compartment surrounding a central lumen (Figure 3H middle, arrows). We observed retention of some hypoblast, though not surrounding the whole epiblast (asterisks). Embryos cryopreserved in 20% DMSO/Sucrose were the most successfully recovered following *in vitro* culture (Figure 3H right). We observed the epiblast (arrows) surrounding the amniotic cavity covered by a fully intact hypoblast (asterisks). Taken together, the optimisation of our cryopreservation protocol to vitrification is also reflected in whole embryo morphology and survival rates post-thawing. These results lay the foundation for a study optimising *in vitro* culture methods for pre- and peri-gastrulation embryos.

### Optimised Cryopreservation and Thaw Protocol

Following these optimisation steps, we developed the final vitrification and thawing protocol which we describe here in detail. Eggs were collected at 0dpo (within 24h of laying) and cleaned first using a dry tissue to remove dirt and then with RNAseZap. Eggs were cut in half and the yolk carefully removed while eggshells were transferred to Tyrode’s solution. Here, the omphalopleure, which lines the inner side of the eggshell was isolated, from which the embryo was cut out of the surrounding tissue using scissors and then cleaned using needles or forceps (Figure 4A).

**Figure 4.**
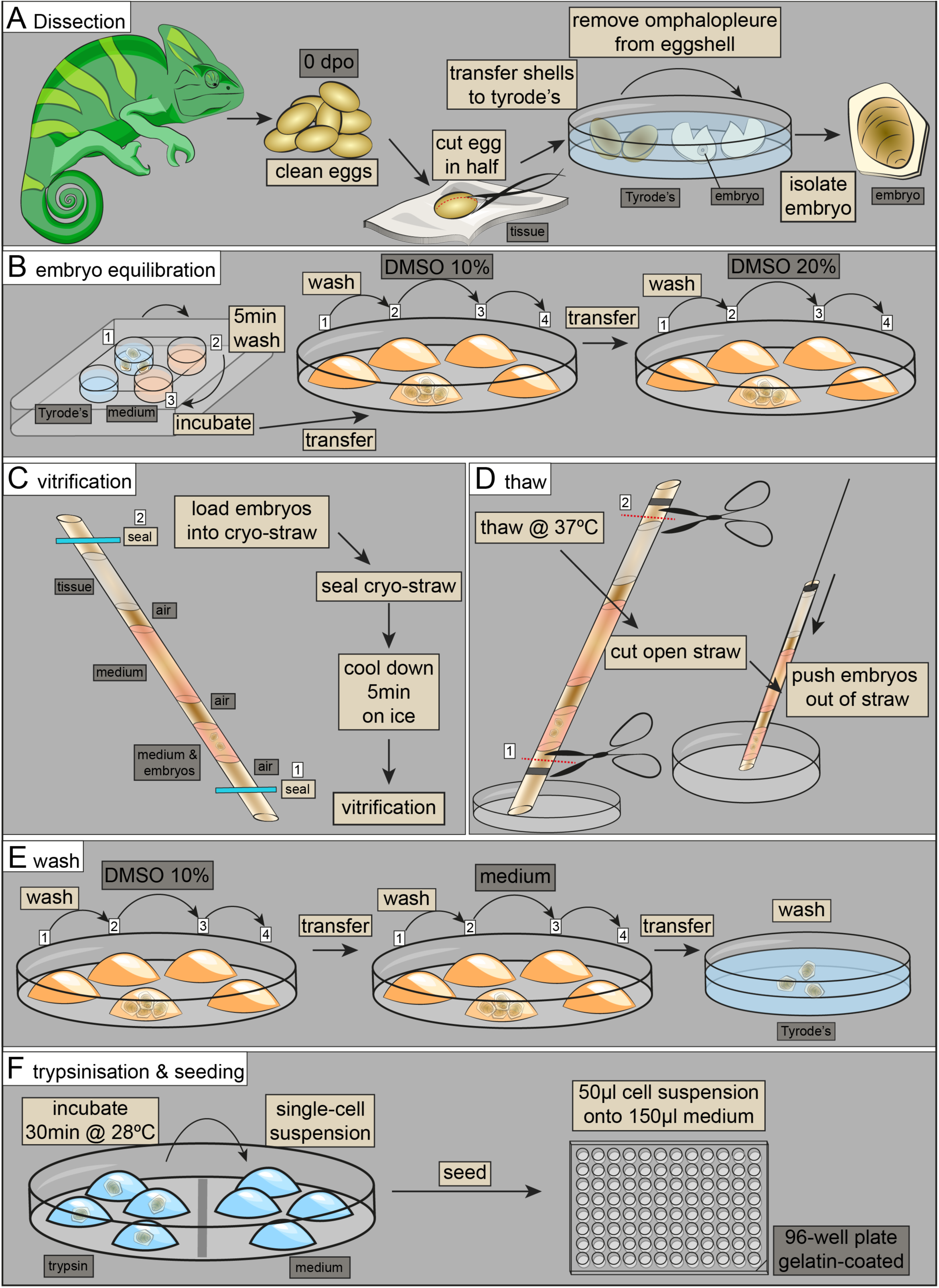
Optimised Cryopreservation and Thaw protocol. **A.** Dissection. 0dpo embryos are first cleaned and then cut in half to remove the yolk. Eggshell halves were transferred to Tyrode’s solution to isolate the omphalopleure, from which the embryo is isolated. **B.** Embryo equilibration. Embryos were collected in Tyrode’s solution and then washed in culture medium. Embryos were then transferred to freezing medium with 10%DMSO. Embryos were washed through 4 drops of 10% DMSO and then transferred into 20% DMSO freezing medium, where they were also washed through 4 drops of 20% DMSO. **C.** Vitrification. Embryos were loaded into a cryo-straw, which is sealed on either side and then equilibrated on ice for 5min before vitrification in LN2 vapour. **D.** Thaw. Straws are thawed in a water bath. Then the seals are cut off and the embryos pushed out of the straws using a small plunger. **E.** Washes. Embryos were washed through 4 drops of 10% DMSO and then transferred to culture medium, where they were also washed through 4 drops. Embryos were then washed in an excess of Tyrode’s solution to remove serum. **F.** Trypsinisation & Seeding. Embryos were transferred into individual drops of trypsin/EDTA and incubated for 30min at 28°C. To stop the reaction, the embryos were transferred to individual drops of medium and suspended to a near-single cell suspension. The suspension was seeded into a gelatin-coated well of a 96-well plate containing 150ul of culture medium.

The embryos were then transferred to culture medium (Shylo et al., 2024) until all the embryos were dissected. To equilibrate the embryos into freezing medium, the embryos were first washed by transfer through 4 drops of 10% DMSO freezing medium (culture medium containing a total of 50% serum (40% chick embryo serum, 10% FBS) and 0.35M Trehalose/0.5M Sucrose) and then incubated in the final drop for 5min before being washed by transfer through 4 drops of 20% DMSO freezing medium, again with incubation in the final drop for 5min (Figure 4B). This ensures optimal and gentle penetration of the cryoprotectant through the entire embryo.

The embryos were then transferred to a cryo-straw (0.5cc) (Figure 4C), which was sealed using a heat sealer. The straw was transferred to ice to cool down for 5min and then vitrified in LN2 vapour for 5min before being submerged in liquid nitrogen. The straws were stored in liquid nitrogen until future use.

To thaw, the cryo-straw was rapidly thawed in a water bath at 37°C. The moment, the ice was thawed, the straw was taken out of the water bath and wiped clean. The seals were cut open carefully as to not lose the embryos, which were then collected in a tissue culture plate using a small plunger (Figure 4D). It is important to prevent air bubbles at this stage and to carefully watch the embryos, which are visible to the bare eye, to ensure they are not lost when being pushing out of the straw.

The embryos were then directly transferred to culture medium containing 10% DMSO and washed 4 times. This is to gently remove the DMSO and not shock the embryos (Figure 4E). The embryos were then transferred to another culture plate with pure culture medium and washed again through 4 drops of medium before being transferred to an excess of Tyrode’s solution to wash out the serum. This step is essential for successful subsequent embryo dissociation.

The embryos were dissociated individually in small drops of trypsin/EDTA for 30min at 28°C and then transferred with minimal trypsin carry over to 50ul drops of culture medium (Figure 4F) to stop the reaction. Through vigorous pipetting, the embryo was broken apart into a near-single cell suspension, and again it is very important to avoid air-bubbles. The cell-suspension was then transferred to a well of a 96-well plate containing 150ul of culture medium which was previously coated with gelatin. Taken together, this presents our optimised and highly versatile cryopreservation and thawing protocol for squamates.

## Discussion

Cryopreservation of embryos is a vital tool for conservation, developmental biology and stem cell research. However, no cryopreservation protocol exists for non-avian reptile embryos. In this study, we initially developed a tissue culture method that supports the culture of cells derived from early gastrulating chameleon embryos at 0dpo. Although the cells did not attach to plastic, all coating methods resulted in cell attachment. We then decided to culture all the cells on gelatin-coated wells, as this is a very common lab resource and our goal was to make our protocol as accessible as possible.

Both vitrification and slow freezing of either single cell suspension or whole embryos yielded surviving cells. To not only serve the goal of stem cell derivation but also enable whole embryo culture for embryo manipulation or spatial analysis methods, we decided to optimise the whole embryo cryopreservation method. Vitrification is more suitable than the more complex process of slow freezing in a fieldwork setting, and we therefore decided to optimise vitrification. Although we optimised this protocol in a lab, we are confident its simplicity can ensure the cryopreservation of specimens in the field. To be able to seed a cell suspension for tissue culture, we dissociated the embryos post thawing using trypsin which is an enzyme that best works at 37°C. However, chameleon cells must be cultured at 28°C. As such, the trypsinisation time, which is commonly between 3-10min for mammalian applications, was prolonged to 30min at 28°C. Over-trypsinisation may affect later cell survival, however generating a cell suspension required considerate effort. We did not succeed in generating a complete single cell suspension, which would have required even longer trypsinisation and much more vigorous pipetting. However, we decided that a near-single cell suspension with small clumps would be sufficient to assess the success of our cryopreservation protocol. To generate a proper cell culture method, requires further optimisation in a future study.

The addition of Trehalose or Sucrose had a marked effect on embryo and single cell survival which may also be achieved by adding a different sugar as an external cryoprotectant. We focused on Trehalose and Sucrose due to their accessibility as common lab chemicals. Our cryopreservation protocol differs with respect to DMSO concentration and the addition of external cryoprotectants when compared to protocols commonly used for mammalian embryo preservation (Dobrinsky, 2002; Khaydukova et al., 2024). The differences may be necessary due to the relatively large size and density of a gastrulating chameleon embryo compared to a mammalian blastocyst. Furthermore, we observed a relatively high variability in total cell numbers from embryo to embryo within a specific cryopreservation method when we optimised the protocol. This may be due to the fact that the chameleons are not inbred such as mouse lines and as such the 0dpo embryo exhibits a natural variability in size.

Nonetheless, our method, which is optimised for early gastrulating chameleon embryos should be adaptable to other squamate embryos. Depending on embryo thickness, a lower DMSO concentration may be more favourable, which would need to be tested. However, our method provides a starting point for researchers aiming to cryopreserve peri-gastrulation squamate embryos in the lab or in the field. By providing a detailed record of our optimisation steps, we present a roadmap for how our protocol may be adapted to different species.

Taken together, our cryopreservation method will not only be highly useful in the hands of conservation scientists to preserve early embryos and the primordial germ cells of endangered species, but could also prove vital for the generation of embryonic stem cells of specimens collected in a field, thereby enabling the molecular study of embryogenesis in the highly evolutionary diverse squamate order.

### Limitation of Study

In this study, we pioneer a cryo-preservation protocol for a peri-gastrulation squamate embryo, the veiled chameleon. In contrast to established amniote model systems, such as mouse or chicken, it is more complex to access 0dpo chameleon embryos due to the fact that a female chameleon lays eggs only every 3-4 months. As such, we had to work with lower specimen numbers (n) than is typically possible with established model organisms such as mouse or chicken. While we recently described pre-oviposition morphogenesis (Weberling et al., 2025) and have annotated the chameleon genome (Shylo et al., 2024), we have not yet been successful in defining tissue specific marker genes or identifying specific antibodies of marker genes, which hinders a definite assessment of the tissue types cryopreserved using our method. The cells grow on gelatin-coated plastic dishes but not on glass which did not allow us to carry out higher resolution imaging to gain more insight into cellular morphology. Polymer coverslips that enable high resolution imaging may be a suitable alternative.

### Contributions

AW conceived of, planned and carried out the project with the help of MD. Cryo-preservation and thawing were performed by AW and MD. NAS provided critical feedback. CMK supported the whole-well imaging of cultured cells and carried out cell number analysis. HW carried out paraffin-sectioning and Histology. RK carried out chameleon breeding and husbandry. SAW provided critical feedback and mentoring. PAT supervised the project. AW wrote the manuscript with the help of MD, NAS, SAW, and PAT.

## Supporting information

combined supplementary Figures

## Acknowledgements

The authors thank all members of the Trainor and Williams lab for valuable discussions and feedback. We thank Dr Connor Ross for critical feedback. Research in the Trainor lab is supported by the Stowers Institute for Medical Research, and N.A.S is supported by a K99/R00 Pathway to Independence award from the National Institute of Child Health and Human Development (HD114881). AW is supported by a postdoctoral research fellowship of All Souls College, University of Oxford.

## Methods

### Data availability

Upon publication, original data provided in this manuscript can be accessed from the Stowers Original Data Repository at https://www.stowers.org/research/publications/libpb-xxxx

### Chameleon Husbandry

The Stowers Institute for Medical Research is an AAALAC accredited institution. Chameleons were housed according to SIMR IACUC approved protocols 2022-147 Lizards and Snakes (Squamata) and 2023-160 Squamate Development as published previously (Baumann and Kupronis, 2021; Diaz et al., 2015). In summary, chameleons were housed in partially screened enclosures measuring about 2’x2’x4’ that were cleaned daily. UV sources and basking opportunities were provided though Mercury vapor and T5 lighting. Each enclosure is set up with a variety of artificial and natural vines, branches and foliage. A variety of insects and vegetables were provided once daily on a rotating basis as food with dietary supplementation given every other feeding. To provide sufficient humidity, misting and whole room humidification occurred several times daily. All chameleons used in this study were housed individually except during documented mating events. Breeding age females were provided lay tubs containing a sand/soil/peat moss mixture. These tubs were misted as needed to maintain appropriate moisture levels. Eggs were removed for artificial incubation within 24h of oviposition (0dpo). No females were sacrificed in this study.

### Embryo Dissection

A clutch of eggs was collected at 0dpo, and wiped with a tissue, to remove large pieces of dirt. As the eggs are under pressure it is essential to be careful and not use too much force. The eggs were then cleaned with RNAzap. At 0dpo, it is impossible to see the embryo by candeling (shining a light through the shell) as is custom for later stages. Instead, the eggs were cut in half on a piece of tissue paper, and the individual shells were placed in Tyrode’s solution. The yolk was discarded by carefully dipping the eggshell halves on the tissue. The embryo is embedded in the omphalopleure, which is attached to inner side of the eggshell. The eggshell was therefore pinched with one set of forceps while a second was used to loosen the omphalopleure from the shell. Once loosened, it can be peeled off. A thick, loose layer of yolk may be on top of the omphalopleure, which is removed by parting the two layers at one section and then pulling them apart using forceps. The embryo is most easily distinguishable from the dorsal side as a darker circle with a light, round structure in the middle – the actual embryo. The omphalopleure around the embryo is cut with forceps and scissors or a needle and then the embryo is cleaned further with needles or forceps, to remove a maximum of surrounding tissue. The embryos were then transferred to a plate of fresh Tyrode’s solution and then culture medium (culture media (DMEM/F-12 (Gibco 10565-018), 10% fetal bovine serum, 15% chicken embryo extract, 1X antibiotic antimycotic solution (Sigma Millipore A5955-100 ML) and 1µg/ml gentamycin solution (VWR 97062-974))(Shylo et al., 2024) or fixated in ice-cold 4% PFA (w/v) in PBS.

### Cryopreservation

#### Slow Freeze Trials

The embryos were equilibrated in freezing medium (10%DMSO, 10% fetal bovine serum, 40% chicken embryo extract, 1X antibiotic antimycotic solution (Sigma Millipore A5955-100 ML) and 1µg/ml gentamycin solution (VWR 97062-974) in DMEM/F-12 (Gibco 10565-018)) for 5min and then moved to a 2ml cryovial. For single cell freezing, the embryos were dissociated in 20ul of Trypsin/EDTA for 30min at 28°C, transferred to 100-200ul droplets of freezing medium without DMSO and separated through vigorous pipetting. The cells were then placed in 500ul of freezing medium in a cryovial. The cryovial was placed on ice for 10min and then moved into a -20°C freezer in a Styrofoam container to control the freezing rate. After 5h, the container was moved to -80°C overnight. The following day, the vials were moved to liquid nitrogen.

#### Vitrification Trials

The embryos were equilibrated in freezing medium by moving them through droplets of freezing medium for 5min. Then, the embryos were loaded onto cryo-straws and sealed using a heat sealer. The straws were placed on ice for 5min and then vitrified in liquid nitrogen vapour before submerging in liquid nitrogen. Single cell suspensions were frozen using the same method but in cryovials. For each condition, 5 embryos were vitrified in each experiment. Following thaw 3 embryos were then suspended and seeded while the other two were fixed and their morphology assessed.

#### Final vitrification protocol

The embryos were stepwise equilibrated in freezing medium, by transferring them into freezing medium (0.5M Trehalose/0.25M Sucrose, 10% FBS, 40% chicken embryo extract in DMEM/F-12 (Gibco 10565-018), 1X antibiotic antimycotic solution (Sigma Millipore A5955-100 ML) and 1µg/ml gentamycin solution (VWR 97062-974))) containing 10% DMSO (v/v) for 5min, after which they were transferred into 20% DMSO (v/v) for 5min at room temperature. Two embryos were loaded per straw in the minimal volume of freezing medium. Following sealing, the straw was moved to ice for 5min to equilibrate and then vitrified in liquid nitrogen vapour before being submerged in liquid nitrogen and transferred to a tank for storage

### Thawing

#### Trials

Cells were thawed in a water bath until only a small clump of ice was visible. 1ml of DMEM/F12 was mixed with the cell suspension and the cells were transferred to a 15ml falcon filled with 10ml of DMEM/F12 to dilute the DMSO. The cells were centrifuged at 0.2rcf at room temperature for 5min. The supernatant was discarded and the cells resuspended in culture medium and seeded for culture.

#### Embryos

The straws were thawed in a water bath, and the embryos were pipetted out of the straws and washed in drops of culture medium to remove the DMSO. The embryos were then individually moved to 20ul droplets of Trypsin/EDTA and trypsinised for 30min at 28°C. The embryos were then transferred to a drop culture medium and separated into a near-single cell suspension through vigorous pipetting. The cell suspension was then transferred to a 96-well plate well filled with 150ul of medium and cultured for 48h.

#### Final protocol

The straws were thawed in a water bath, and the embryos pushed out of the straws and transferred into droplets of culture medium containing 10% DMSO. The embryos were washed 4 times and then moved into drops of culture medium, where they were washed 4 more times before being transferred into a 4-well plate well filled with 1ml of culture medium. The embryos were then washed in Tyrode’s solution to remove the serum and individually placed in 20-40ul droplets of trypsin/EDTA for 30min at 28°C. The embryos were then transferred to 50ul droplets of culture medium, carrying over the minimum of trypsin. The embryos were broken apart into a near-single cell suspension by vigorous pipetting while minimising the generation of air bubbles. The cell suspension was then seeded into a 96-well plate and cultured for 48h.

### Embryo Culture

The embryos were cultured at 5% CO2 in a humidified atmosphere at 28°C. The embryos were placed in chameleon fibroblast culture medium which consisted of 10% fetal bovine serum and 15% chicken embryo extract in DMEM/F-12 (Gibco 10565-018), with 1X antibiotic antimycotic solution (Sigma Millipore A5955-100 ML) and 1µg/ml gentamycin solution (VWR 97062-974))(Shylo et al., 2024). 1-3 embryos were cultured in one well of a 96-well plate in 150ul culture medium for 24h.

### Tissue Culture

Cells were cultured at 5% CO2 in a humidified atmosphere at 28°C. Wells of a 96-well plate were coated with gelatine for 1h at 28°C. The gelatin was removed and after about 5min of air drying, 150ul of medium was added to the wells. The embryos were individually trypsinised for 30min in 20ul of trypsin and then moved to a droplet of 50ul. Each embryo was separated through vigorous pipetting, and the cell suspension was seeded in a 96-well plate well. The cells were assessed after 24h of culture and fixed after 48h of culture. For tissue culture trials, the wells were alternatively coated with L-glutamine, which followed the same procedure as gelatin coating. For matrigel embedding, matrigel was thawed on ice and an embryo was moved from Trypsin/EDTA through a drop of medium and then separated following trypsinisation in 30ul of ice-cold matrigel using a cold tip. The matrigel solution was then pipetted into a 96-plate well. To solidify the matrigel, the plate was moved to 28°C for 5-10min and then covered with 150ul of medium. To seed cells on a matrigel bed, 30ul of matrigel were used to cover the well, incubated at 28°C for 5-10min to solidify, and then covered with medium that contained the cell suspension. The medium contained 5% (v/v) matrigel.

### Immunofluorescence staining

All incubation and washing steps were performed on a rocker unless stated otherwise. The embryos were fixed in 4% PFA (in PBS) at 4°C overnight, washed 3x for 10min in PBS and permeabilised for 30min in permeabilisation buffer (0.3M TritonX100, 0.1M glycine in PBS). Then, the embryos were rinsed in PBST (PBS, 0.1% Triton X100) and blocked for 1h at room temperature in blocking buffer (0.1% Triton X-100, 1% v/v donkey serum in PBS). The cells were fixed in 4% PFA at room temperature for 20min, washed 3x in PBS for 5min, and permeabilised for 20min. After a brief rinse in PBST, the cells were blocked for 1h if required, followed by primary antibody incubation. If the cells were only stained for DAPI and Phalloidin, all steps from blocking to the washes after the primary antibody incubation were skipped.

Primary antibodies were applied to the embryos and cells overnight at 4°C in blocking buffer. The embryos were washed 3x for 10min, whereas cells were washed 3x for 5min. Secondary antibodies were applied to the embryos and cells at room temperature, which were kept in the dark on a rocker. The embryos were incubated for 3-4h, whereas the cells were incubated for 2h. The cells were washed 4x 10min in PBST and imaged in the 5^th^ wash. This minimised background fluorescence. The embryos were washed 3x 10min and then equilibrated for 30min each in 25%, 50% glycerol before being mounted in Vectashield.

Primary Antibodies used: goat-anti-Brachyury (1:200), rabbit-anti-phosphoHistoneH3 (1:200). Secondary antibodies used: donkey-anti-goat AF568 (1:500), donkey-anti-rabbit AF568 (1:500). Fluorescent stains: DAPI (1:500), Phalloidin-AF488 (1:500).

### Imaging

The embryos were imaged on a Zeiss LSM 980 or on a Leica SP8. All replicates for an individual experiment were imaged on the same microscope. The cells were imaged on a Leica SP8 for qualitative image analysis. For each well which contained cells originating from an individual embryo, two areas were imaged at a 10x air objective on a Leica SP8. Nucleus size was measured following the StarDist nuclear detection plugin (Schmidt et al., 2018). Only nuclei between 60-350um size were quantified to remove apoptotic nuclei or large, merged multi-nucleic structures. Segmented areas that had a near to background mean gray value as well as areas that had near-maximum mean gray value were excluded from further analysis.

The whole wells were imaged for quantitative analysis using a Nikon Ti2 equipped with a Yokagawa CSU W1 Spinning Disk. DAPI was imaged with a 405 laser 430-480 nm emission filter. Z-stack images were taken in a 5x5 tile with a 10x NA0.45 objective, then stitched and maximum projected in Fiji. Cell counting was completed in Python 3.14 with Cellpose3 using a diameter of 17 and flow threshold of 0.6 to segment and count nuclei.

### Histology

Embryos were washed 3x in PBS for 5min and then dehydrated into EtOH in a stepwise manner 30/50/70% EtOH (v/v) for 5min each at room temperature. To lightly dye the embryos in order to facilitate correct orientation during embedding, a couple of drops of Eosin were added to the 70% EtOH. The embryos were then processed for paraffin sectioning (Milestone, Pathos Delta Microwave Tissue Processor) according to the following protocol without pressure: 70% EtOH for 4 min at room temperature, 100% EtOH for 5 min at 37°C, Isopropyl Alcohol for 5 min at 45°C, Paraffin for 15 min at 62°C. Once processing was completed, the embryos were embedded in paraffin wax (Cancer Diagnostics, PureAffin® R56) and sectioned at a thickness of 5 µm on a microtome (Leica RM2255) along their anterior-posterior axis. Automated HCE staining was carried out using a stainer (DP360, Dakewe (Shenzen) Medical Equipment Co.) with ST Infinity HCE Reagents (Leica Biosystems Cat. 3801698). Slides were mounted with Cytoseal 60 mounting media (VWR, 48212-187).

The embryos were imaged using an Olympus Slide Scanner on a 20x objective. For each section, a z-stack of 10 slices 1um apart was imaged to ensure capturing of the correct focal plane due to the individual sections being too small for the auto-focus function of the slide scanner. For display and analysis, the z-slice most in focus was chosen.

## Figure legends

**Figure S1: Freezing trial**

**A.** Further examples of the results of freezing trials. DAPI (cyan) and F-Actin (magenta) staining. Boxes indicate high magnification insets. **B.** Immunofluorescence staining of embryos vitrified, trypsinised and seeded onto gelatin-coated wells in medium containing matrigel. Fixation 48h post seeding. DAPI (cyan) and F-Actin (magenta) staining. Boxes indicate high magnification insets. Example images from 3 biological replicates. All scale bars 100um.

**Figure S2: Cryopreservation optimisation assay**

**A/B.** Additional examples of the optimisation assay shown in Figure 2B. Maximum intensity projections of cells cultured for 48h following thawing and single cell suspension. (A) Embryo vitrified in 10% DMSO (+/-) Trehalose/Sucrose. (B) Embryo vitrified in 20% DMSO (+/-) Trehalose/Sucrose. DAPI (cyan) and F-Actin (magenta) staining. Squares indicate regions of zoom-in.

**Figure S3: brightfield assessment of whole embryo culture**

**A.** Brightfield images of 0dpo embryos. ventral side. **B.** Brightfield images of embryos following thawing. Top row: embryos frozen in 20% DMSO plus Sucrose. Bottom row: embryos frozen in 20% DMSO plus Trehalose. **C.** Brightfield images of embryos cultured for 24h post thaw. Top row: embryos frozen in 20% DMSO plus Sucrose. Bottom row: embryos frozen in 20% DMSO plus Trehalose. All scale bars 100um.

