## Supplementary figures and images for "A versatile cryopreservation method for peri-gastrulation squamate embryos optimised using the veiled chameleon (*C. calyptratus*)"

### combined supplementary Figures

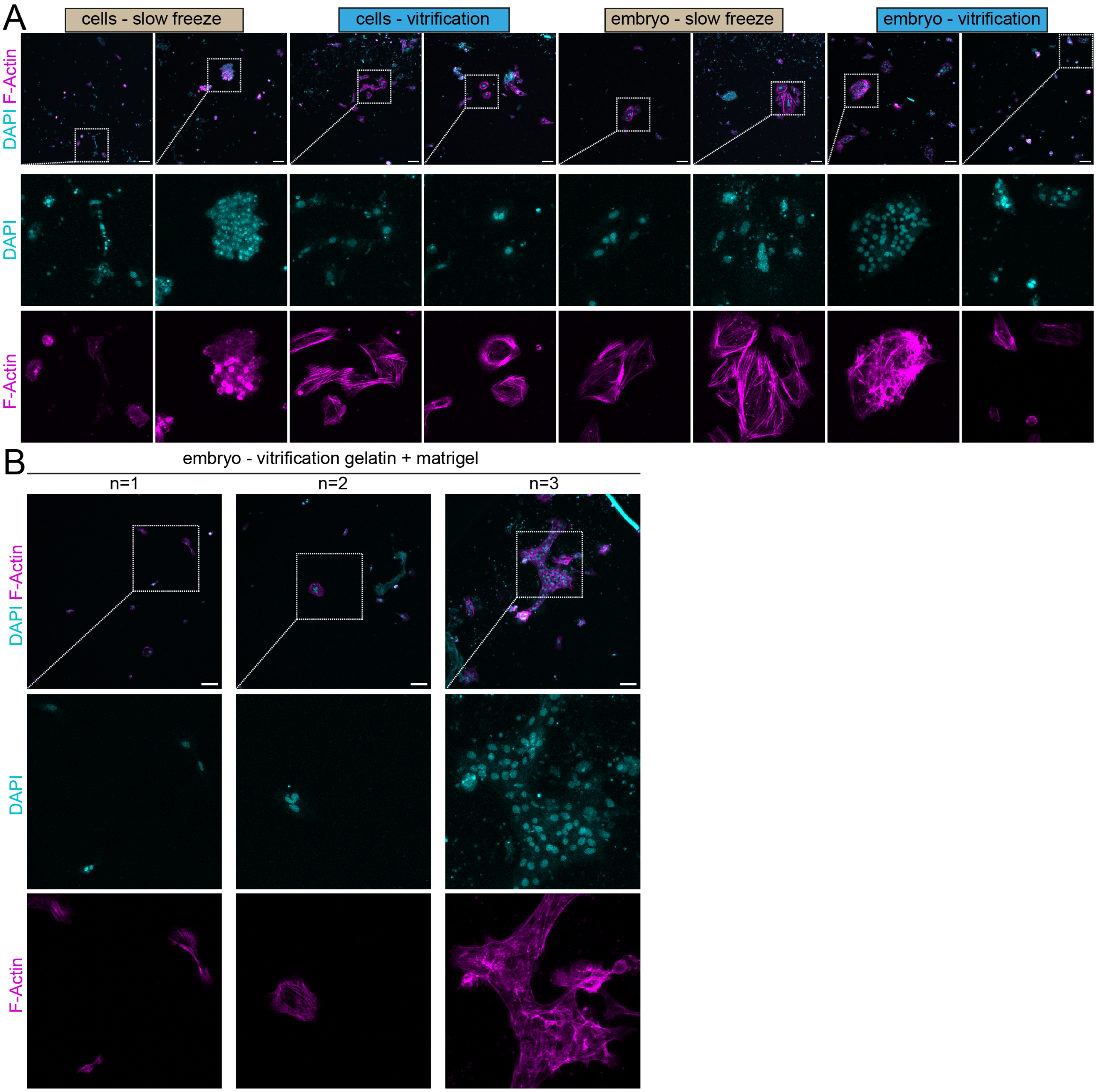

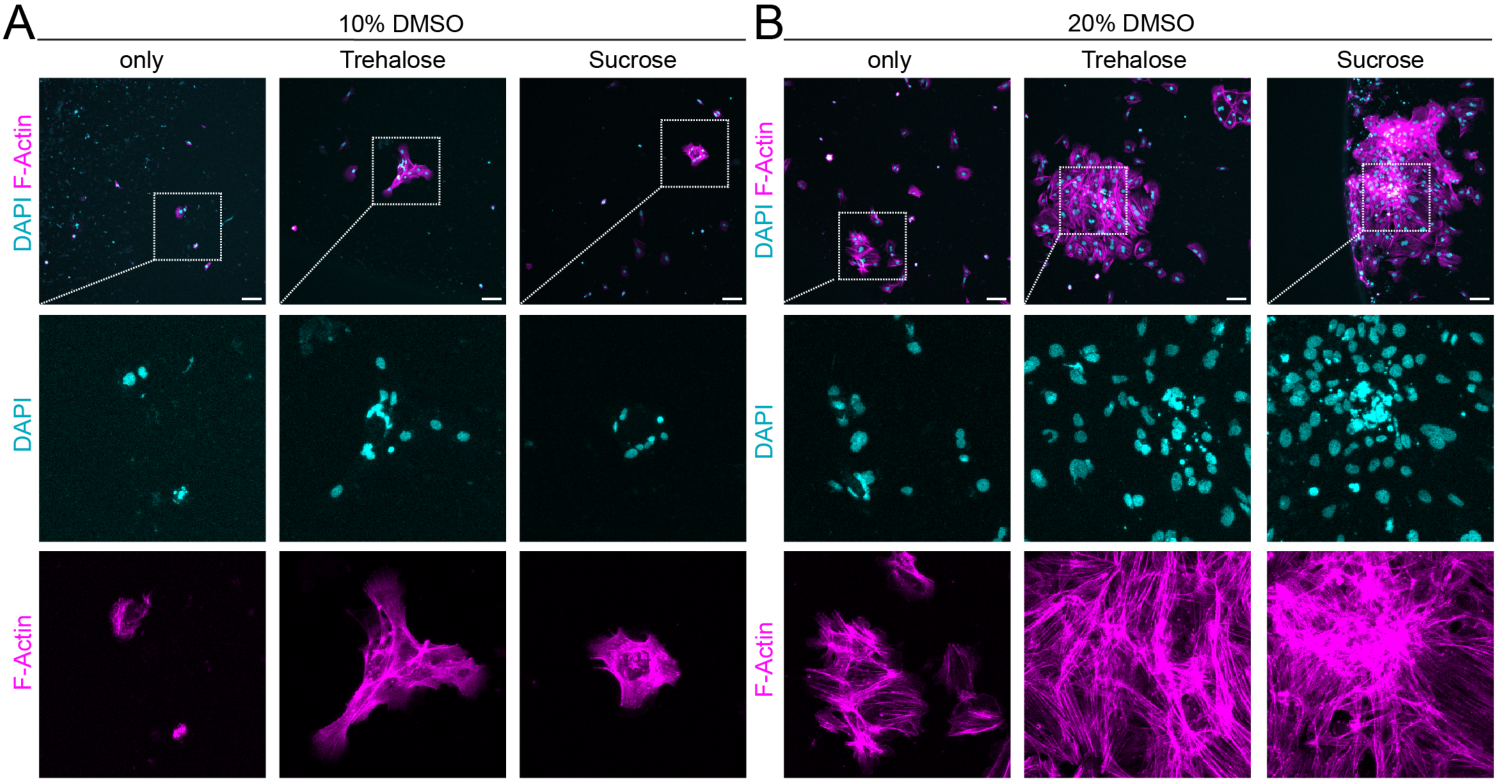

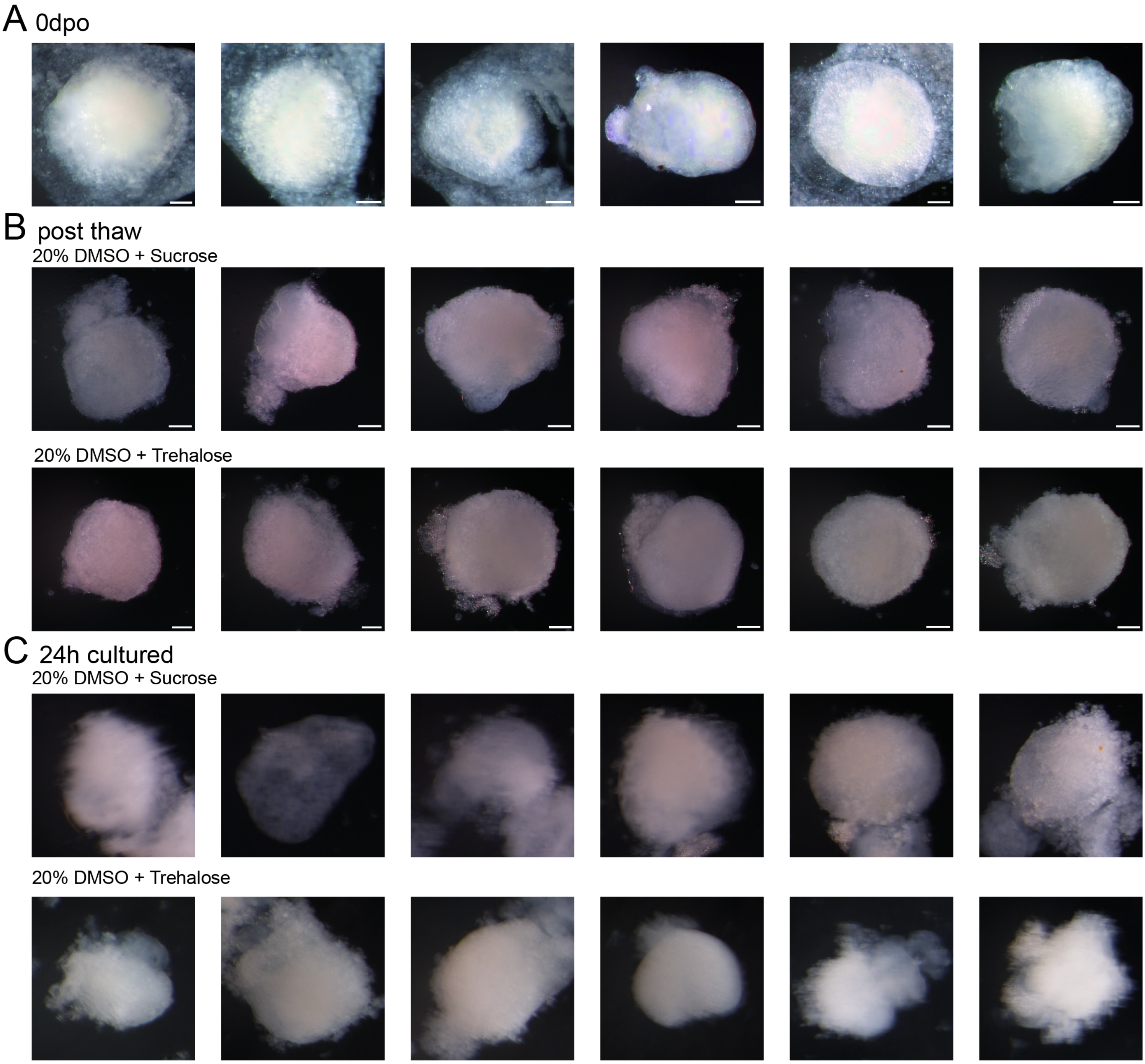
